# A heuristic method to identify runs of homozygosity associated with reduced performance in livestock

**DOI:** 10.1101/131706

**Authors:** J.T. Howard, F. Tiezzi, Y. Huang, K.A. Gray, C. Maltecca

## Abstract

While for the most part genome-wide metrics are currently employed in managing livestock inbreeding, genomic data offer, in principle, the ability to identify functional inbreeding. Here we present a heuristic method to identify haplotypes contained within a run of homozygosity (ROH) associated with reduced performance. Results are presented for simulated and swine data. The algorithm comprises 3 steps. Step 1 scans the genome based on marker windows of decreasing size and identifies ROH genotypes associated with an unfavorable phenotype. Within this stage, multiple aggregation steps reduce the haplotype to the smallest possible length. In step 2, the resulting regions are formally tested for significance with the use of a linear mixed model. Lastly, step 3 removes nested windows. The effect of the unfavorable haplotypes identified and their associated haplotype probabilities for a progeny of a given mating pair or an individual can be used to generate an inbreeding load matrix (**ILM**). Diagonals of **ILM** characterize the functional inbreeding load of individual (IIL). We estimated the accuracy of predicting the phenotype based on ILL. We further compared the significance of the regression coefficient for IIL on phenotypes to genome-wide inbreeding metrics. We tested the algorithm using simulated scenarios (n =12) combining different levels of linkage disequilibrium (LD) and number of loci impacting a quantitative trait. Additionally, we investigated 9 traits from two maternal purebred swine lines. In simulated data, as the LD in the population increased the algorithm identified a greater proportion of the true unfavorable ROH effects. For example, the proportion of highly unfavorable true ROH effects identified raised from 32 to 41 % for the low to the high LD scenario. In both simulated and real data the haplotypes identified were contained within a much larger ROH (9.12-12.1 Mb). The IIL prediction accuracy was greater than zero across all scenarios for simulated data (high LD scenario mean (95% confidence interval): 0.49 (0.47-0.52)) and for nearly all swine traits (mean ± SD: 0.17±0.10). On average across simulated and swine datasets the IIL regression coefficient was more closely related to progeny performance than any genome-wide inbreeding metric. A heuristic method was developed that identified ROH genotypes with reduced performance and characterized the combined effects of ROH genotypes within and across individuals.

## INTRODUCTION

The implementation of routine genotyping within livestock breeding populations has become a common practice and is used as a tool to make more effective selection decisions in swine breeding companies (Knol et al., 2016). Previous research has highlighted the advantages of genomic relationships compared to pedigree-based information to obtain more precise estimates of the genetic merit and homozygosity of an individual (Knol et al., 2016; Lopes et al., 2013). Also, genomic information allows genome-wide inbreeding estimates to be supplemented by characterizing the impact of homozygosity for specific genomic regions. Elevated levels of homozygosity result in a reduction in phenotypic performance, referred to as inbreeding depression (Falconer and Mackay, 1996). The identification of region-specific stretches allows breeders to manage inbreeding more effectively since the impact of homozygosity for a trait can vary across the genome. The estimation of dominance effects to identify unfavorable regions of the genome have been utilized in the past (Lopes et al., 2016; Xiang et al., 2016), but lacks power for low frequency mutations and doesn’t consider that whole segments of the genome are passed from parent to offspring. To overcome these limitations, regions of the genome in a continuous run of homozygosity (ROH) have been proposed to investigate homozygous segments that arose due to past inbreeding (Howard et al., 2015; Saura et al., 2015). Previous research has investigated the phenotypic effect of a region being in an ROH (Pryce et al., 2014; Howard et al., 2015). The previous methods did not directly identify the unique ROH genotype that gave rise to the reduced phenotypic performance. Within this paper, we have attempted to close this gap by developing a heuristic algorithm to identify unfavorable haplotypes contained within ROH across the genome. The method was tested on simulated as well as real data.

## MATERIALS AND METHODS

No animal care approval was required for this work since all genotypes and records came from data that were available from previous studies. The manuscript will be split in two sections. In the first section, we will provide an overview of the algorithm along with methods to summarize the number of unfavorable haplotypes shared within and across individuals. In the second part of the paper we will employ simulated and swine data sets to summarize three major results: (1) how effective the algorithm is at identifying unfavorable haplotypes, (2) the length of ROH the unfavorable haplotype tags, (3) the relationship of the aggregate effect of unfavorable haplotypes carried by an individual with its phenotype and genetic value.

### Description of the algorithm

A pictorial overview of the algorithm is displayed in **Figure 1**. The method follows three steps. The first step scans the genome to identify ROH genotypes that result in an unfavorable change in the phenotype of interest. The genotypes utilized in the algorithm are coded as 0 for the homozygote, 2 for the alternative homozygote and 1 for the heterozygote. Step 1 begins at the first SNP of a chromosome by constructing a window of a predetermined number of SNP (default = 60). Within a window, the mean phenotype is tabulated for each unique ROH genotype and any genotype that is not in a ROH is aggregated into a category referred to as nonROH. Furthermore, any ROH genotype below a user defined frequency (default = 0.0075) is removed from the ROH genotype list and placed in the nonROH category. If the phenotypic mean for a ROH genotype is below/above a user defined value (discussed below), the window is stored. Next, the window is shifted forward by one SNP and the previous process is repeated. Once the entire chromosome has been scanned, windows containing the same set of animals and representing the same ROH genotype except for the first and last SNP are aggregated (i.e. **Figure 1**: Step 1b). Since recombination does not occur within a given region for the individuals of the same ROH genotype, ROH genotypes that are combined in this step contain the same amount of information. Following the aggregation of nested windows, the window length is reduced by “n” SNP (default = 5) and the previous steps are repeated for the new window size. The window size is reduced by “n” until a minimum window size is reached (default = 20). Once the minimum window size is reached, the process of scanning for unfavorable ROH genotypes is complete for a chromosome. For windows that contain the same set of animals and are nested within each other (i.e. **Figure 1**: Step 1d), the shortest window is kept for further analysis. The aggregation steps (i.e. Step 1b and 1d) trap the ROH genotypes across individuals that have the same core ROH genotype of the smallest possible length. No information is lost within the aggregation steps as each ROH genotype belongs to the same set of individuals, yet the step significantly reduces the number of windows tested in following steps. The core ROH genotype is now expected to serve as a tag for the full ROH segment observed in an individual, which may differ across subjects due to recombination occurring at different locations across subjects.

**Figure 1.**
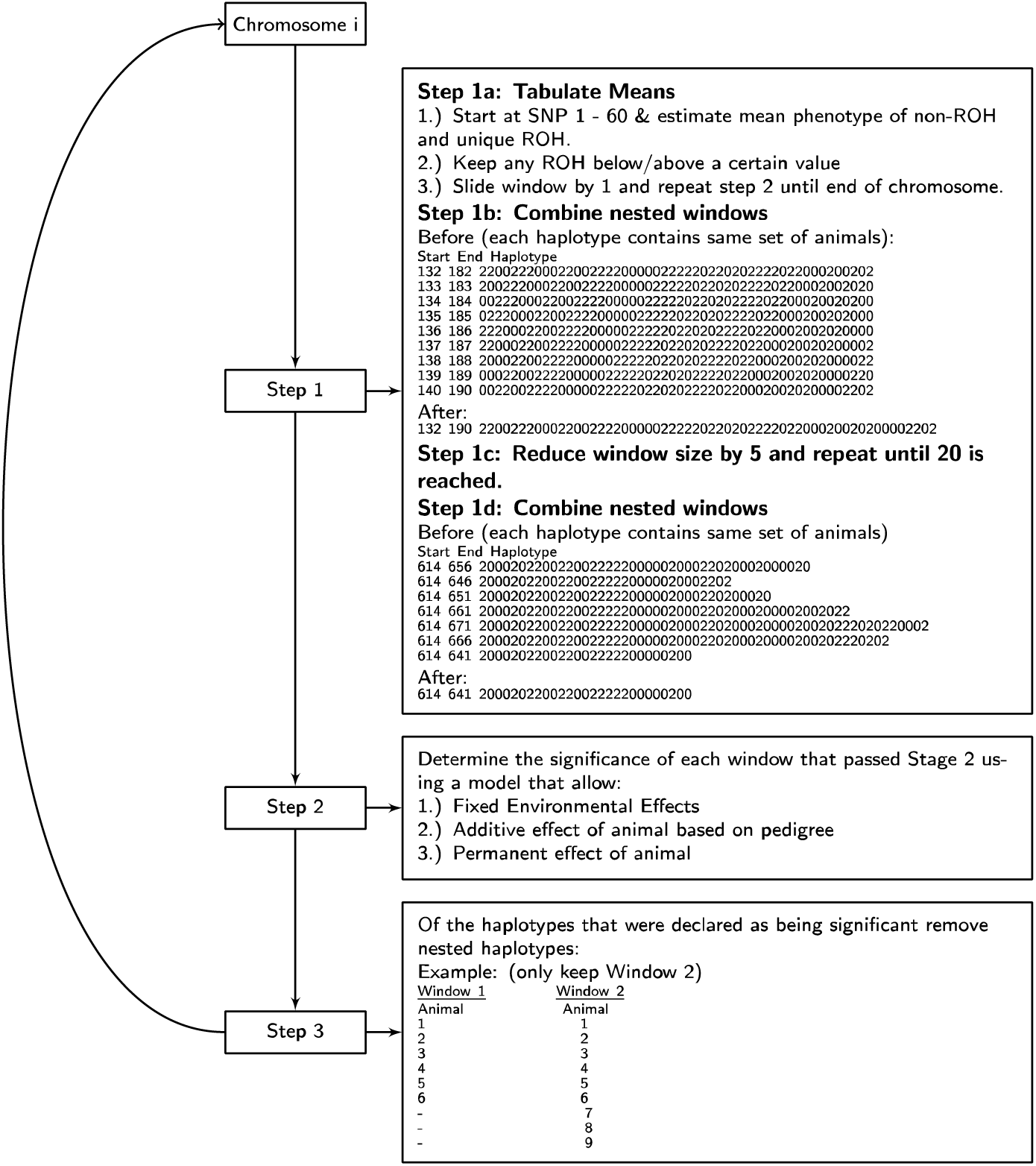
Overview of algorithm that identifies unfavorable haplotypes.

Any window remaining after Step 1 is subsequenly tested for significance using a standard mixed model that accounts for the environment, additive genetic and permanent environment effect of an individual, plus any number of fixed effects. A description of the full model for each window is outlined below:

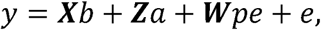

where *y* is the trait of interest, *b* is a vector of fixed effects, *a* is a vector of random additive genetic effects, *pe* is a vector of random permanent environmental effects, *e* is a vector of random residuals and **X**, **Z**, and **W** are incidence matrices relating *b*, *a* and *pe* with *y*, respectively. The fixed effects can include any environmental classification or covariate effect along with the effect of ROH genotype (i.e. unique ROH genotype and nonROH) for a given window. The random additive genetic effect is assumed 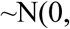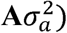, with **A** representing the additive relationship matrix derived from a pedigree (Henderson, 1976). The random permanent environmental and residual effects are assumed 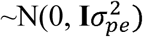 and 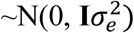, respectively, with **I** being an identity matrix. Variance components of the current implementation are assumed fixed across windows based on the null model of no ROH effect (i.e. no ROH genotype in the model). For each window, solutions are obtained via the Cholesky decomposition of the left-hand side (**LHS**). Given the solutions for each window, a contrast between each unique ROH genotype versus nonROH and the associated t-statistic are obtained. The contrasts 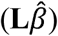 are obtained following Welham et al. (2004) and Gilmour et al. (2004). The t-statistic is generated based on the following formula:

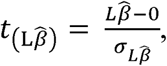

where 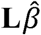 refers to the estimated contrast and 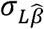 refers to the standard error of the contrast, calculated as **L**(**LHS^-1^**)**L’**. The hypothesis test is one-sided and the direction of the test is dependent on the direction of the unfavorable phenotype. Under this parameterization, the genotypes not in an ROH are assumed normal compared to individuals that have the ROH genotype. This aligns with the partial dominance hypothesis, which is thought to account for the majority of inbreeding depression observed in populations (Simmons and Crow, 1977; Charlesworth and Charlesworth, 1987). Any contrast that passes the user-defined significance threshold is kept and moved onto the final window reduction step which resolves nested windows (i.e. **Figure 1**: Step 3).

The algorithm presented was developed in C++11. The source code and compiled executable files for Linux operating systems are available at “https://github.com/jeremyhoward”. The primary option the user controls is the cutoff value for the mean phenotype for a given ROH genotype that is considered unfavorable in Step1a. The user can specify a cutoff value based on prior knowledge of what is considered an unfavorable phenotype or generate an empirical t-statistic distribution from the data to declare a cutoff value. The latter option is conducted by randomly specifying a chromosome, window length and start position and estimating the significance value for ROH genotypes within the window. All one-sided t-statistics are stored. Across samples, the mean phenotype for t-statistics with a significance ranging from 0.10 and 0.05 is chosen as the cutoff value.

The haplotypes identified can be utilized in a variety of ways, but two are investigated in the current study. The first application is to apply the algorithm across economically important phenotypes and identify haplotypes having an unfavorable effect across multiple traits. Regions with consistent unfavorable effect across multiple traits should have a high probability of being sensitive to inbreeding and thus result in a reduction in the overall fitness and vigor of an individual. The second application is to generate a matrix aiming at characterizing the decrease in the trait of interest across all unfavorable haplotypes, herein referred to as the inbreeding load matrix (**ILM**). Its calculation follows the method outlined by Cole (2015). In order to implement an **ILM**, the genotype phase needs to be known. This matrix then can be utilized in mating designs to minimize the probability of progeny containing the unfavorable haplotype(s) for a single trait or across multiple traits. The diagonals of the matrix, referred to as individual inbreeding load (IIL), represent an individual’s decrease in the phenotypic performance(s) due to inbreeding, while off-diagonals represent the decrease in the trait(s) of the progeny given the mating of the two (potential) parents. It should be noted that in this implementation the algorithm does not run all haplotypes across the genome simultaneously. As a result, any observed ROH genotype for an individual might contain multiple significant unfavorable haplotypes. Therefore multiple tag haplotypes identified by the algorithm could be counted as different in an individual, when in fact they are tagging the same observed haplotype. Within the current study, when multiple haplotypes tagged the same observed ROH genotype, only the haplotype with the highest significance value and resulting in the largest number of haplotypes observed across individuals was retained. For the i^th^ row and j^th^ column of **ILM**, the following formula was utilized to calculate the value:

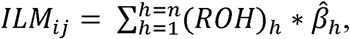

where *n* is the number of unfavorable haplotypes that remained after eliminating haplotypes that were not observed or removed to avoid double counting. The *ROH*_*h*_ refers to the probability of generating a ROH for haplotype_h_ and *β*_*h*_ is the effect of the ROH genotype estimated from Step 2 of the algorithm. The probability values for the diagonals elements of the ILM include 0.25 (haplotype carrier) or 1.0 (haplotype in ROH). The probability values for the off-diagonal elements include 0.25 (mating of haplotype carriers), 0.5 (mating of haplotype carrier and ROH genotype) or 1.0 (both parents have ROH genotype). The **ILM** values range from 0 (i.e. no unfavorable haplotypes) to any value in the unfavorable direction.

### Summary of metrics utilized to test the algorithm using simulated and swine data

Simulated datasets, where the true genetic signal is known, were employed to determine how effective the algorithm was at identifying true negative ROH regions as well as to characterize the relationship between IIL and the true aggregate genotypic value of individuals. The length of ROH the unfavorable haplotype tagged was determined across both simulated and swine datasets in order to ensure that long stretches of ROH were represented. Long ROH stretches have a higher probability of being true IBD segments as a result of recent inbreeding, compared to shorter ones. The relationship between IIL and the phenotype was summarized based either on a) IIL accuracy of predicting the phenotype or b) the significance of the regression coefficient when IIL was included as a fixed covariate in a mixed linear model. The latter relationship was generated under the premise that the management of inbreeding is traditionally done by minimizing parental coancestries using genome-wide inbreeding metrics. Therefore, the significance (i.e. *–log p-value*) of the regression coefficient from traditionally utilized genome-wide inbreeding metrics was compared with the IIL value. Lastly, IIL was benchmarked across simulated and swine datasets with estimates of the genetic value based on a whole genome regression model. This was conducted to generate a reference comparison on the prediction accuracy for a given trait based on traditionally utilized genome-wide modeling techniques. It is important to note that the genetic signal from IIL encompasses only unfavorable effects resulting from long IBD segments. As a result, a comparison of the prediction accuracies between the two metrics needs not to be interpreted as an exercise of ranking the predictive ability of the two metrics, (ILL value would by construction only capture a subset of the overall genetic signal) but rather to determine the relationship between complementary metrics, in order to allow their integration.

### Simulated Data

Multiple scenarios were simulated to determine the frequency of unfavorable haplotypes being identified by the algorithm. Simulation was carried out using Geno-Diver (Howard et al. In Press), a combined coalescence and forward in time simulation software. We hypothesized that the amount of short-range LD existing in the genome impacts how well the algorithm can identify unfavorable haplotypes. Four scenarios of increasing levels of short-range LD in the historical population were generated as outlined in **Figure S1** and will be referred to as the *low*, *low-medium*, *medium-high* and *high*, respectively. For each LD scenario, different genetic architectures were simulated with 250, 500 or 1000 QTL spread equally across 5 chromosomes. The combination of variable LD and QTL parameters produced 12 different scenarios. Each scenario was replicated 25 times.

Within each LD setting, SNP sequence data for 4000 base haplotypes across 5 chromosomes, each with a length of 150 Megabases, were simulated by internally calling MaCS (Chen et al., 2009) within the Geno-Diver software. Scenarios with the same LD parameter were initialized using the same set of sequence data to limit the computational time and variability across replicates due to historical sequence information. Following the generation of sequence data, QTL were randomly placed along the genome and a SNP panel with neutral markers was created. A total of 4,000 markers (20,000 genome-wide) were utilized within each chromosome. This marker density was chosen to generate a density within each chromosome that is similar to a medium density marker array such as the Illumina PorcineSNP60K (Illumina Inc., San Diego, CA). Across all scenarios, the minimum minor allele frequency was set at 0.10 and 0.015 for markers and QTL, respectively.

For each QTL the additive effect (*a*) of a QTL, defined as half the difference in genotypic value between the homozygote genotypes (Falconer and Mackay, 1996), was sampled from a gamma distribution (shape = 0.4; scale = 1.66) with an equal chance of being positive or negative. The dominance effect (*d*) of a QTL, defined as the deviation of the genotypic value of the heterozygote from the mean of the genotypic values of the two homozygotes (Falconer and Mackay, 1996), was generated similarly to Wellmann & Bennewitz (2012). First, the degree of dominance (*h*) at QTL_i_ was sampled from a normal distribution (mean = 0.1; variance = 0.04) and then the dominance effect at QTL_i_ was calculated as d_i_ = *h*_i_|*a*_i_|, where |*a*_i_| is the absolute value of the additive effect. Across all scenarios, the additive and dominance effects were scaled to generate a narrow and broad sense heritability (H^2^) of 0.35 and 0.40, respectively. The normal distribution parameters used to generate the degree of dominance were utilized to create a trait that displayed directional dominance along with a majority of the loci displaying partial dominance. Phenotypes were simulated by adding a residual value, generated from a normal distribution (mean = 0, variance = (1- H^2^)), to the genotypic value for each animal. Summary statistics on the QTL architecture and genetic diversity of the 12 scenarios is outlined in **Table S1.**

After the founder population and genetic architecture of the trait was generated a selection scenario mimicking a livestock population was undertaken for ten generations. A population consisting of 50 males and 600 females was utilized, with a replacement rate of 20% for both males and females. Progeny with a high estimated breeding value (EBV) were selected to serve as parents for the next generation and EBV were generated from an animal model based on pedigree information. A low phenotypic value represented the unfavorable direction for the simulated trait in this case. Animals were mated at random and one progeny was produced for each mating pair. Progeny born from generation 7 to 9 served as the training population to identify unfavorable haplotypes and progeny from generation 10 served as the validation population. The model utilized to identify unfavorable haplotypes in the simulation data set did not have a permanent environmental effect since individuals only had 1 observation and the only fixed effect was the overall mean. The starting window size was set at 60 and was reduced by 5 until a window size of 20 SNP was reached. Different SNP window sizes were investigated based on the density simulated. Similar results were found in terms of the regions identified, associated effects and its relationship with the phenotype (data not shown). The suggestive phenotypic cutoff in step 1 was declared by randomly sampling 1000 windows to generate the empirical t-statistic distribution.

To investigate the proportion of true negative ROH effects the algorithm captured within each replicate, the true effect for any ROH with a length greater than 1 Mb was calculated. A length of 1 Mb was chosen to provide a range of possible ROH lengths captured by the algorithm. The true negative and positive ROH effects were split into quantiles of decreasing and increasing effects, respectively. The algorithm only tests for the unfavorable direction and therefore the percentage of true ROH effects the algorithm identified is expected to be higher in the negative compared to the positive direction. Lastly, using the same 1 Mb ROH cutoff, statistics on the length of ROH the algorithm identified (or missed) were calculated.

Within each replicate, the ILL was estimated based on haplotypes identified in the training population for individuals in the validation population. The correlation between IIL and the true genotypic value (TGV), true breeding value (TBV) and true dominance deviation (TDD) was also estimated. Additionally, the significance (i.e. *–log p-value*) of IIL or a genome-wide metric when included as a fixed covariate effect was estimated for the validation population. The ILL or genome-wide metric was included as a fixed covariate in the similar model (i.e. no ROH effect included in model) that was used to identify haplotypes in the training population. Three genome-wide inbreeding metrics were used as comparison including pedigree inbreeding (Henderson, 1976), diagonals of the SNP-by-SNP relationship matrix (**SNPRM**; VanRaden, 2008) or proportion of the markers that were homozygous.

To explore the predictive ability of IIL compared to estimates of the genetic value utilizing whole genome regression models, a Bayesian Ridge Regression (BRR) analysis was conducted that included the additive and dominance effect for each SNP. The same training and validation generations that were utilized previously were also used in the BRR analysis. Marker effects were estimated using the ‘BGLR’ package in R (Perez and de los Campos, 2014). A total of 55,000 iterations were run with the first 5,000 discarded as burn-in and a thinning rate of 5. Across individuals, the estimated breeding value (EBV), dominance deviation (EDD) and genotypic value (EGV) were generated by multiplying the estimated effect by the associated genotype and summing across all markers. The prediction accuracy for either IIL or EGV was determined in the validation population based on the correlation between phenotype and EGV or IIL, respectively. It was standardized by dividing by the square root of the heritability estimated in the training generation for each replicate (Legarra et al., 2008; Wolc et al., 2011). Correlations between IIL and the EBV, EDD or EGV were also estimated.

### Swine Data

Phenotypic and genotypic data from two maternal purebred nucleus selection lines were obtained from Smithfield Premium Genetics (Rose Hill, NC). In order to determine the algorithm’s behavior across different genetic architectures, multiple traits were investigated including litter size, litter viability and growth rate. Individuals with genotype information from Large White (LW, n = 6,750) and Landrace (LR, n = 5,010) were utilized. Animals born before 2012 were used as a training population and animals born on 2013 were used as a validation population and the number of animals across traits is outlined in **Table 1**. A complete description of the genotype quality control is outlined in Howard et al. (2016). Briefly, genotype data was derived from the Illumina PorcineSNP60K BeadChip (Illumina Inc., San Diego, CA) and the GGP-Porcine (GeneSeek Inc., a Neogen Co., Lincoln, NE). Multiple quality control measures were conducted before imputing and phasing missing and low-density to medium-density genotypes using Beagle (Version 3; Browning & Browning 2007). After quality control and discarding SNP that were poorly imputed, a total of 39,671 and 41,489 autosomal SNP for LW and LR remained, respectively.

**Table 1.** The model utilized, estimated genetic parameters and the number of animals across traits for the Landrace (LR) and Large White (LW) population.

| Breed | Trait<sup>1</sup> | Fixed Effects<sup>1</sup> | Genetic Parameters<sup>2</sup> |  | Animals (Records) |  |
| --- | --- | --- | --- | --- | --- | --- |
| | | | $h^2$ | $r^2$ | Training | Validation |
| LR | NBA | Parity, CG | 0.082 | 0.149 | 4,005 (9,416) | 1,005 (1,639) |
|  | TNB | Parity, CG | 0.082 | 0.149 | 4,003 (9,302) | 998 (1,621) |
|  | PD | Parity, CG | 0.085 | 0.156 | 4,003 (9,293) | 998 (1,621) |
|  | LBW | Parity, CG, NBA | 0.231 | 0.284 | 3,985 (8,648) | 988 (1,586) |
|  | PWM | Parity, CG, Pig24hrs | 0.115 | 0.131 | 3,504 (6,724) | 870 (1,330) |
|  | NW | Parity, CG, Pig24hrs | 0.102 | 0.120 | 3,465 (6,600) | 845 (1,280) |
|  | NWBW | Parity, CG, Pig24hrs, NW | 0.144 | 0.211 | 3,465 (6,600) | 845 (1,280) |
|  | Weight | CG, Sex, Age | 0.271 | - | 4,386 (4,386) | 993 (993) |
|  | ADG | CG, Sex, Age | 0.271 | - | 4,386 (4,386) | 993 (993) |
| LW | NBA | Parity, CG | 0.115 | 0.150 | 5,518 (15,014) | 1,232 (2,271) |
|  | TNB | Parity, CG | 0.098 | 0.142 | 5,513 (14,673) | 1,228 (2,262) |
|  | PD | Parity, CG | 0.086 | 0.133 | 5,513 (14,664) | 1,228 (2,262) |
|  | LBW | Parity, CG, NBA | 0.241 | 0.307 | 5,487 (13,581) | 1,188 (2,155) |
|  | PWM | Parity, CG, Pig24hrs | 0.074 | 0.168 | 4,966 (10,464) | 1,102 (1,824) |
|  | NW | Parity, CG, Pig24hrs | 0.053 | 0.115 | 4,901 (10,259) | 1,054 (1,716) |
|  | NWBW | Parity, CG, Pig24hrs, NW | 0.144 | 0.203 | 4,901 (10,259) | 1,054 (1,716) |
|  | Weight | CG, Sex, Age | 0.292 | - | 5,576 (5,576) | 1,197 (1,197) |
|  | ADG | CG, Sex, Age | 0.291 | - | 5,576 (5,576) | 1,197 (1,197) |

<sup>1</sup> NBA = number born alive; TNB = total number born; PD = proportion born dead; LBW
= average litter birth weight; PWM = pre-weaning mortality; NW = number weaned;
NWBW = average litter wean weight; Weight = weight at off-test; ADG = average daily
body weight gain from birth to off-test; CG = contemporary group based on farm, year
and season; Pig24hrs = pigs in the litter after the 24 hour cutoff; Age refers to off-test
age.
<sup>2</sup> $h^2$ refers to the narrow sense heritability; $r^2$ refers to the repeatability.

Seven litter size and mortality traits including number born alive (NBA), total number born (TNB), proportion born dead (PD), average litter birth weight (LBW), preweaning mortality (PWM), number weaned (NW) and average litter wean weight (NWBW) were employed in the analysis. The TNB phenotype included NBA, stillborn and mummified piglets. The PD dead was calculated as 1 - (NBA/TNB). The LBW was calculated as the mean weight of the number of live piglets at processing, which occurred within 48 h from birth. Traits that were recorded after birth, including PWM, NW and NWBW, are impacted by the degree of cross-fostering. Cross-fostering in the current data was similar to previous estimates by Putz et al. (2015) in a related population. To minimize the effect of cross-fostering only litters having more than 75 % of the birth sow piglets were utilized in the analysis. After the data edit, 98.0 and 97.7 % of the piglets were nursed by their original birth sow for LW and LR, respectively. The PWM mortality phenotype was calculated as the number of piglets that died after 24 hours including pigs euthanized at weaning divided by the total number of pigs in the litter after the 24-hour cutoff. The NWBW was calculated as the average weight of the number of piglets weaned. All reproductive traits were evaluated as a trait of the biological dam. The fixed effects utilized for each trait are outlined in **Table 1.** A random additive genetic and permanent environmental effect of the dam were included in the analysis (i.e. similar to Model 1 described in the section outlining the algorithm).

Two production traits were investigated: body weight at off-test and average body weight gain from birth to off-test (i.e. body weight at off-test / age at off-test). Production traits were evaluated as a trait of the animal. Since animals only have one observation the permanent environmental effect was in this case excluded. The fixed effects utilized for each trait are described in **Table 1**. Across both reproductive and production traits, the contemporary group (CG) was comprised of farm, year and season and any animal that was within a CG smaller than 5 was removed from the analysis.

Summary statistics on the length of ROH and the unfavorable haplotypes captured were generated. Prediction accuracy for IIL was compared with a whole genome regression BRR model. For all 9 traits, yield deviations were constructed for each trait based on the fixed effects outlined in **Table 1**. For the reproductive traits, an animal may have multiple observations and therefore average yield deviations were used and the residuals for a given observation in the BRR analysis was weighted according to Garrick et al. (2009). The formula used to calculate the weight was:

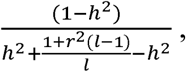

where *h*^2^ refers to the heritability, *r*^2^ refers to the repeatability and *l* refers to the number of records. The values used for *h*^2^ and *r*^2^ are outlined in **Table 1** across all nine traits. Utilizing ‘BGLR’ (Perez and de los Campos, 2014), a total of 155,000 iterations were run with the first 5,000 discarded as burn-in and a thinning rate of 5. Again, for each trait the EGV values were predicted. The accuracy of predicting the phenotype utilizing IIL compared to a whole-genome regression model and the relationship between the two were investigated. The prediction accuracy for either IIL or EGV across the nine traits was determined in the validation population. The prediction accuracy was calculated as the correlation between the EGV or IIL and average yield deviation. It was standardized by dividing the square root of the heritability for each trait. The significance of the ILL or genome-wide regression coefficient was estimated as outlined previously.

Lastly, the correlation between the diagonal and the off-diagonal elements of **ILM** across traits and with pedigree- and genomic-based relationship matrices were estimated. Understanding the correlation between **ILM** across traits is important when **ILM** is used to minimize inbreeding depression across all traits in a breeding objective. Any change in the off-diagonal **ILM** value for one trait should ideally result in a favorable or negligible change in the off-diagonal **ILM** value for other traits. **ILM** was compared to three relationship matrices including pedigree-based (**A;** Henderson, 1976), **SNPRM** and a ROH-based relationship matrix with a 5 Mb cutoff (**ROH5RM**; Howard et al. 2016). To determine the sensitivity of fixing the variance components based on the null model of no ROH effect, the ASReml program (Gilmour et al., 2009), which re-estimates variance components for each window was utilized across breed and traits for the windows that were deemed significant by the algorithm.

## RESULTS

### Simulated Data

A summary of how effectively the algorithm identified true negative and positive ROH effects across different percentiles is outlined in **Panel 1 of Figure 2**. Since the algorithm only tests for the unfavorable direction, the percentage of true ROH effects the method identifies is expected to be greater than zero in the negative direction and zero in the positive direction. As illustrated in **Panel 1 of Figure 2**, as the true negative unfavorable ROH effect got larger, a greater proportion of unfavorable ROH genotypes was identified by the algorithm. It should be noted that, averaged across all scenarios, the frequency of highly unfavorable ROH effects was small (1.8 %) compared to the total number of true negative ROH effects. The frequency of incorrectly identified positive ROH effects (i.e. false-positives) by the algorithm remained relatively flat across all percentiles and was on average (95% confidence interval (**CI**)) 9.4 (8.7-10.1) percent across all scenarios. As the LD in the population increased and became similar to that of most livestock populations, the algorithm was more effective at identifying unfavorable haplotypes and had a lower false-positive rate. For example, for true ROH effects with the largest negative effect (i.e. less than the 0.05 percentile), the algorithm identified on average (95% CI) 32.1 (28.5-35.7) and 41.2 (36.7-45.9) percent of the total true negative ROH effects across the three QTL scenarios for the low and high LD scenarios, respectively. Conversely, for incorrectly identified true ROH effects (i.e. estimated to be negative, but had a true positive effect) with the largest positive effect (i.e. greater than the.95 percentile), the algorithm identified on average (95% CI) 15.3 (13.9-16.7) and 9.6 (8.2-10.9) percent of the total true ROH effects across the three QTL scenarios for the low and high LD scenario, respectively.

**Figure 2.**
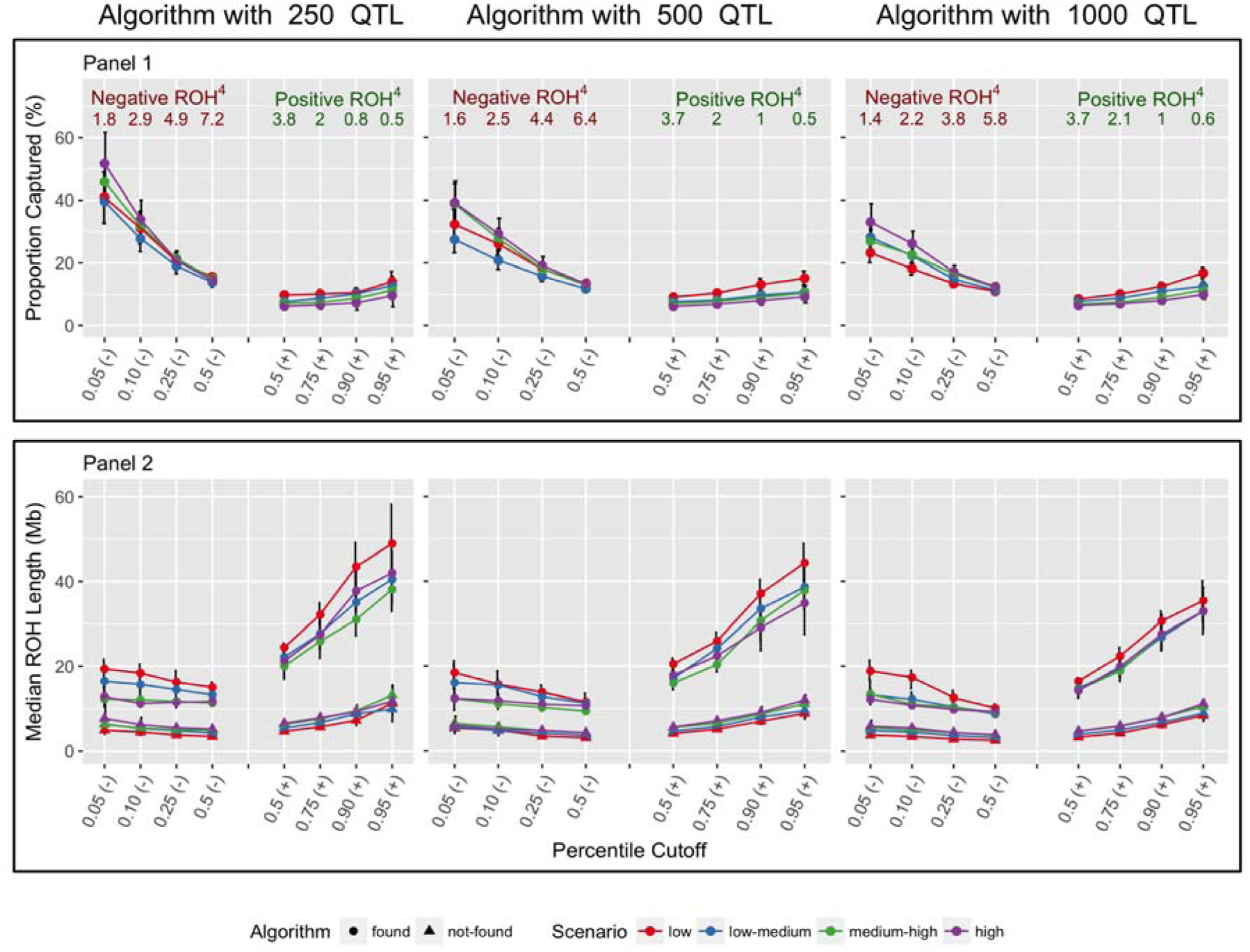
Summary^1^ of the proportion of runs of homozygosity (ROH) of at least 1 Megabase the algorithm captures (Panel 1) and the length of the ROH the haplotype tags (Panel 2) across simulation scenarios^2^ by percentile class^3^ and whether the algorithm identified the haplotype.

Summary statistics on the length of ROH of at least 1 Mb tagged by the unfavorable haplotype is outlined in **Panel 2 of Figure 2**. We report the median in this case, rather the mean, since the distribution of the length of ROH containing a tag haplotype has a heavy tail and thus the latter parameter is heavily influenced by extreme values. The length of ROH tagged by the identified haplotypes for the medium-high and high LD scenarios was similar across negative percentiles and QTL scenarios with a median (1^st^ quartile – 3^rd^ quartile) of 12.15 (10.07-13.41). The haplotypes identified for the low and low-med LD scenarios across negative percentiles and QTL tagged longer ROH stretches with a median length of 15.77 (12.23-18.64). The results show how the core unfavorable haplotype identified by the algorithm, which had a median length of 7.0 kilobases (kb) across scenarios, in reality serves as a proxy for a much larger observed ROH segment. The length of unfavorable ROH that the algorithm missed was made of considerably smaller ROH (median (1^st^ quartile – 3^rd^ quartile): 5.26 (4.06-5.81) Mb) and was again similar across negative percentiles and scenarios. For the incorrectly identified true positive ROH effects, it should be noted that the length of ROH captured by the haplotype gets longer proportional to the true ROH effect. Thus, in general, falsely identified ROH regions were in our analysis characterized by being locally negative around the identified unfavorable haplotype. Yet as a result of being part of an extremely large ROH, positive QTL effects contained in the long ROH genotype made the overall effect positive.

The relationship of IIL with the true genetic signal, the predictive ability of ILL compared to whole genome regression values and the significance of IIL or genome-wide inbreeding regression coefficients are outlined in **Figure 3**. **Panel 1 of Figure 3** describes the correlation between IIL with TGV, TBV, and TDD. Across all QTL scenarios, the correlation increased as the LD increased for all parameters except for TDD. Averaged (95% CI) across QTL scenarios the correlation between IIL and the TGV for the low and high LD scenario was 0.31 (0.29-0.32) and 0.44 (0.42-0.45), respectively. The correlation between IIL and TBV were similar to the correlations between IIL and TGV. The average (95% CI) correlation between IIL and TDD was 0.002 (-0.01-0.01) for all scenarios, which was not unexpected, given the fact that the ROH effects are a function of the alternative homozygote genotypes and not heterozygous genotypes.

**Figure 3.**
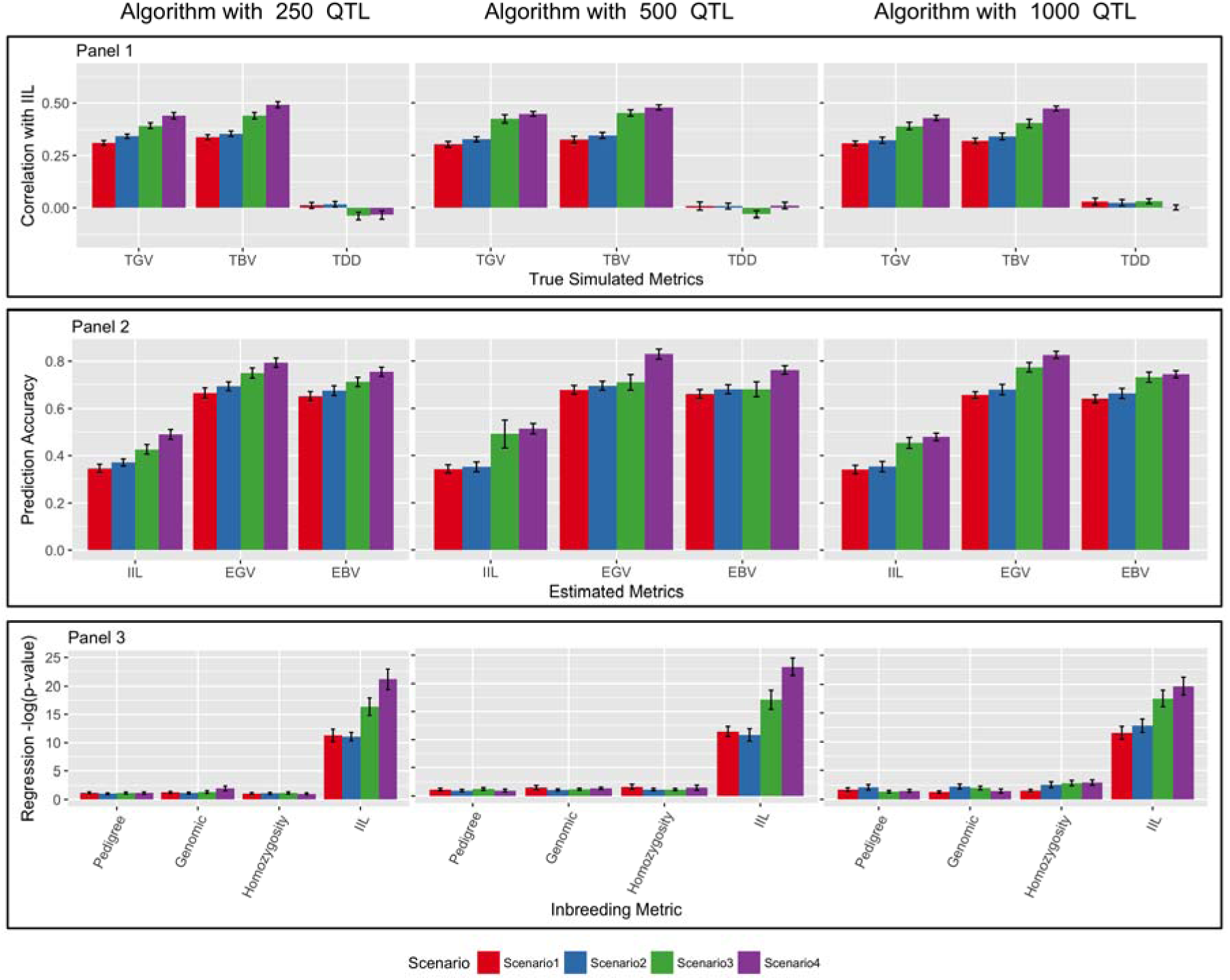
Summary (mean ± CI)^1^ of the correlation between individual inbreeding load (IIL) and the true genetic signal (Panel 1)^2^, prediction accuracy for IIL and genotypic estimates Bayesian ridge regression (Panel 2)^3^ and significance of the regression coefficient based on either a genome wide inbreeding metrics or IIL (Panel 3)^4^ across simulation scenarios^5^.

**Panel 2 and 3 of Figure 3** summarize the effectiveness of using the IIL algorithm based on its predictive ability or as a tool to minimize the frequency of unfavorable haplotypes in the progeny. As outlined in **Panel 2 of Figure 3**, the correlation between IIL and the phenotype increased as the level of LD increased in the population. Averaged (95% CI) across QTL scenarios, the prediction accuracy of IIL was 0.34 (0.32-0.36) and 0.49 (0.47-0.52) for the low and high LD scenarios, respectively. Similar trends of increasing prediction accuracy as the LD in a population increased were seen, as expected, for the whole genome prediction values and minor differences were found between the prediction accuracy for EGV and EBV. Averaged (95% CI) across QTL scenarios, the prediction accuracy of EGV was 0.66 (0.64-0.67) and 0.82 (0.79-0.84) for the low and high LD scenarios, respectively. These results are not unexpected since the algorithm only utilizes haplotypes that have an unfavorable effect contained within ROH stretches and favorable haplotypes are not included in IIL. The correlations between IIL and values from the whole-genome regression model are outlined in **Figure S2 Panel 1**. Averaged (95% CI) across scenarios the correlation between IIL and EGV was 0.50 (0.49-0.50) and in general as the LD increased so did the correlation.

The last summary statistic is outlined in **Panel 3 of Figure 3** and outlines the significance of the regression coefficient based on either genome wide inbreeding metrics or IIL. Across all genome-wide inbreeding metrics the *–log p-value* was similar across all LD scenarios, and the significance increased proportionally to the number of QTL. For example, averaged (95% CI) across scenarios and genome-wide inbreeding metrics, the average *– log p-value* were 1.12 (1.01-1.23) and 1.68 (1.49-1.87) for the scenarios with 250 and 1000 QTL, respectively. The *– log p-value* for the IIL metric across all scenarios was in all cases greater and increased as the LD in the population increased. Under the high LD scenario the average (95% CI) *–log p-value* for the IIL metric across QTL scenarios was 21.24 (19.37-23.10), corresponding to a nominal p-value of 5.96*e*^*-10*^.

In summary, the simulation results highlight that the algorithm identified on average 41 % of the highly unfavorable (i.e. 0.05 percentile) ROH effects across the QTL scenarios and under the high LD scenario. Moreover, the unfavorable haplotypes were effective at tagging a significantly larger ROH region. Under the high LD scenario, which closely resembles most livestock situations, the ROH that the haplotype tagged had a median length of 12.1 Mb. When combining all unfavorable haplotypes based on their probability of occurring and the effect of the haplotype being in a ROH, a moderate prediction accuracy was achieved. Furthermore, the correlation between ILL and the EGV was moderate and more importantly less than unity in all cases. Therefore, a combination of ILL and a genome-wide genetic value would allow for two animals with similar genetic values but differ in the number of unfavorable haplotypes contained within long ROH to be distinguished.

### Swine Data

To determine whether similar results were found with real data and to investigate its effectiveness across multiple traits, the algorithm was tested with two swine commercial maternal lines. The significance of the regression coefficient and the predictive ability of ILL compared to genome-wide inbreeding metrics is presented in **Table 2**. Across both breeds and for the majority of traits except for NBA and TNB in LR, ILL had a prediction accuracy greater than 0. Averaged (± SD) across traits within a breed, the average prediction accuracy was 0.15 (± 0.13) and 0.20 (± 0.04) for LR and LW, respectively. Similar to the simulation, the whole genome regression based EGV resulted in higher prediction accuracies compared to ILL across all traits and breed. The prediction accuracy averaged (± SD) across traits within a breed was 0.48 (±0.10) and 0.49 (±0.17) for LR and LW, respectively. Both prediction accuracies were lower than what was achieved in the simulation, given the lower heritability for most of the traits and the simplified assumptions employed in the simulation. The correlations between IIL and values from the whole-genome regression model are outlined in the bottom of **Figure S2 Panel 2**. Averaged (± SD) across traits within a breed the correlations between the IIL and EGV were 0.31 (± 0.13) and 0.32 (± 0.06) for LR and LW, respectively. A positive correlation (Averaged ± SD: LR = 0.07 ± 0.06; LW = 0.15 ± 0.07) was estimated between ILL and EDD.

**Table 2.** The significance of inbreeding regression coefficient across multiple inbreeding metrics and the prediction accuracy of the individual inbreeding load (IIL) and estimated genetic value (EGV) from whole genome Bayesian Ridge Regression across traits for Landrace (LR) and Large White (LW) populations.

| Breed | Trait<sup>1</sup> | Regression on Adjusted Phenotype $-\log(P\text{-value})^2$ | | | | Prediction Accuracy | |
| --- | --- | --- | --- | --- | --- | --- | --- |
|  |  | Pedigree Inbreeding | Genomic Inbreeding | Proportion Homozygous | IIL | EGV | IIL |
| LR | NBA | 0.66 | 0.49 | 0.69 | 0.00 | 0.44 | -0.05 |
|  | TNB | 0.87 | 0.68 | 0.23 | 0.53 | 0.42 | -0.09 |
|  | PD | 0.15 | 0.00 | 1.42 | 1.94 | 0.53 | 0.15 |
|  | LBW | 0.45 | 2.24 | 1.34 | 15.24 | 0.65 | 0.33 |
|  | PWM | 0.29 | 1.54 | 0.8 | 2.94 | 0.40 | 0.17 |
|  | WN | 0.21 | 1.01 | 0.49 | 4.00 | 0.42 | 0.22 |
|  | WNBW | 2.14 | 0.66 | 4.36 | 2.51 | 0.64 | 0.22 |
|  | Weight | 0.53 | 4.52 | 2.93 | 3.24 | 0.40 | 0.17 |
|  | ADG | 0.58 | 4.54 | 2.84 | 4.00 | 0.39 | 0.18 |
|  | Average | 0.65 | 1.74 | 1.68 | 3.82 | 0.48 | 0.15 |
| LW | NBA | 0.62 | 0.15 | 0.12 | 4.14 | 0.45 | 0.24 |
|  | TNB | 0.98 | 0.40 | 0.21 | 2.56 | 0.42 | 0.19 |
|  | PD | 0.54 | 1.35 | 1.97 | 1.94 | 0.38 | 0.13 |
|  | LBW | 2.18 | 1.11 | 4.07 | 10.25 | 0.77 | 0.24 |
|  | PWM | 2.11 | 0.33 | 1.37 | 2.09 | 0.36 | 0.18 |
|  | WN | 1.71 | 0.12 | 0.00 | 2.91 | 0.29 | 0.25 |
|  | WNBW | 0.00 | 0.78 | 2.56 | 3.46 | 0.79 | 0.17 |
|  | Weight | 0.69 | 5.70 | 6.14 | 5.99 | 0.48 | 0.18 |
|  | ADG | 0.73 | 5.76 | 6.09 | 4.77 | 0.48 | 0.17 |
|  | Average | 1.06 | 1.74 | 2.50 | 4.23 | 0.49 | 0.20 |

<sup>1</sup>NBA = number born alive; TNB = total number born; PD = proportion born dead; LBW = average litter birth weight (LBW); PWM = pre-weaning mortality; NW = number weaned; WNBW = average litter wean weight; Weight = weight at off-test; ADG = average daily body weight gain from birth to off-test.
<sup>2</sup> As a reference a p-value of 0.10, 0.05 and 0.01 is equivalent to a negative log p-value of 2.30, 3.0 and 4.6, respectively.

Also, outlined in **Table 2** is the *–log p-value* of the regression coefficient when genome-wide inbreeding or ILL values were included in the model. Averaged across traits within a breed, the IIL regression coefficient resulted in a higher *–log p-value* (i.e. lower p-value) across both breeds compared to any genome-wide inbreeding metric, while the pedigree based inbreeding metric had the lowest *–log p-value*. Out of the 9 traits, the regression coefficient was trending towards significance (P-value < 0.10) for 6 and 7 out of the 9 traits for LR and LW, respectively. Alternatively, the regression coefficient for the genome-wide metrics for LR (LW), was trending toward significance for 3 (4), 2 (2) and 0 (0) of the 9 for the proportion of the genome homozygous, diagonals of **SNPRM** or pedigree-based inbreeding, respectively. Thus, in our results ILL was the parameter that more closely aligned with the identification of functional inbreeding. It should be noted that, no single parameter had a consistently higher *–log p-value* across traits so that a combination of genome-wide inbreeding metric based on genomic information and the IIL value would likely be optimal in breeding applications.

An ideogram of regions of the genome where an unfavorable haplotype was identified by the algorithm across the 9 traits for the two lines is depicted in **Figure S3** and **S4**, respectively. Regions of the genome where long unfavorable stretches of homozygosity were observed across multiple traits/line. Conversely, other regions did not appear to harbor unfavorable stretches of homozygosity. The number of regions that have an unfavorable effect across at least 4 of the 9 traits is outlined in **Table 3** and placed into categories based on the relationship between the traits. A summary of the regions and the least square mean difference between an animal in an ROH versus nonROH across both breeds is outlined in **Table S2.** A total of 4 and 13 regions were found that had at least one production and reproduction trait affected by a tag haplotype in LR and LW, respectively. A total of 3 regions across both breeds were associated only with reproductive traits. Summary statistics on the median ROH length that the unfavorable haplotype tagged and the average frequency of the ROH genotype across traits and breeds is outlined in **Table S3**. The average median length of the unfavorable haplotype across trait and breeds was 1.56 and 1.54 Mb for LR and LW, respectively. Similarly to what found in simulated data, the unfavorable haplotype tagged a larger ROH of 9.55 and 9.12 Mb averaged across traits within LR and LW, respectively, corresponding to (averaged across traits) 172 and 156 SNP for LR and LW, respectively.

**Table 3.** Summary of the number of haplotypes that displayed unfavorable effects across multiple (i.e. > 4) traits for Landrace (LR) and Large White (LW) populations.

| Breed | Type of Trait<sup>1</sup> | Number of Haplotypes |
| --- | --- | --- |
| LR | Production and Reproduction | 4 |
|  | Reproduction | 3 |
| LW | Production and Reproduction | 13 |
|  | Reproduction | 3 |

<sup>1</sup> The type of trait refers a Production trait (i.e. weight at off-test; average daily body weight gain from birth to off-test) or a Reproductive Trait (i.e. number born alive; total number born; proportion born dead; average litter birth weight; pre-weaning mortality; number weaned; average litter wean weight).

Lastly, the correlation between the diagonals and off-diagonals of **ILM** for each trait and genome-wide relationship matrices is presented in **Figure S5** and **S6** for LR and LW, respectively. As shown by the lower diagonal of each matrix, correlations between the off-diagonal elements of the ILM across all traits and genome-wide relationships are all favorably correlation. Any change in the off-diagonal **ILM** value for one trait would result in a similar (in the favorable direction) or negligible change in other traits. The average off-diagonal elements across traits for LR had an absolute correlation of 0.23, 0.28 and 0.34 for A, SNPRM and ROH5RM, respectively. Slightly lower correlations were found for LW and averaged across traits the absolute correlation was 0.14, 0.20 and 0.21 for A, SNPRM and ROH5RM, respectively. In general, the correlations between the IIL values across traits and genome-wide inbreeding metrics were similar to the off-diagonals and the majority of them were in the same direction. In some instances, the correlations between the values were antagonistic, for example LR between the SNPRM and the IIL values across all traits, although the correlations between the genome-wide inbreeding metrics were much lower compared to the off-diagonal elements.

## DISCUSSION

The objective of this study was to implement a strategy to identify haplotypes within long ROH that tag an IBD segment due to recent inbreeding. Haplotypes within ROH were targeted since previous results via simulation by Keller et al. (2011) have shown that ROH based genome-wide inbreeding metrics have a higher association with the recessive mutation load compared to pedigree or SNP-by-SNP based inbreeding metrics. The rationale behind the algorithm proposed stems from previous research investigating the phenotypic effect of a region being in an ROH (Pryce et al., 2014; Howard et al., 2015; Saura et al., 2015). One of the major pitfalls of previously utilized methods is that they assume that any ROH genotype within a region of interest has an unfavorable effect, which is most likely not the case. Instead, the unfavorable effect is likely due to a single unique ROH genotype with the remaining ones resulting in no unfavorable effect. Thus, the necessity of identifying unique ROH genotypes associated with an unfavorable phenotype. The primary outcome of the proposed algorithm is a list of unfavorable haplotypes. Multiple algorithms already exist to manage unfavorable mutations or haplotypes within breeding programs so that the ones identified by the algorithm could be easily incorporated into previously developed pipelines (Kinghorn, 2011; Cole, 2015).

Within the algorithm, multiple aggregation steps are implemented to confine the unfavorable haplotype to the core of the observed ROH genotype in a way that is consistent across individuals. As result of the aggregation step, each haplotype serves as a tag for a much larger ROH segment. In this regard, the data presented confirm that the aggregation steps are successful in identifying tag haplotypes contained within a much larger ROH genotype. Across both swine breeds and in the simulated data set, the median length of the ROH the haplotype tagged was greater than 9 Mb and the tag haplotype was around 1 Mb. Furthermore, simulation results highlighted that the true ROH effects that were not identified were shorter ROH (5.26 Mb) compared to the ones that were identified (13.96). The ability to capture short IBD regions depends on the marker density as described by Ferenčaković et al. (2013) and the marker density utilized in the current study might not be sufficient to capture these short IBD regions effectively. The impact of the density was not investigated here to limit the number of scenarios generated, yet its impact should be considered in the future. Lastly, the simulation highlighted how in some cases the algorithm incorrectly identified true positive ROH effects that were characterized as being much longer compared to correctly identified negative ROH effects. The distribution of the length of ROH has a heavy tail and therefore the frequency of long ROH is low, but they do exist within the genome across individuals. These incorrectly identified true positive ROH regions were locally negative around a tag unfavorable haplotype, but being in longer than average ROH their combined effect was ultimately positive.

We investigated the ability of the algorithm to identify unfavorable haplotypes and their potential use. The frequency at which ROH occurs within the genome had a large impact on the ability of the algorithm to identify unfavorable haplotypes. Medium-high and high LD scenarios have LD patterns similar to those observed in livestock species. Under these premises the algorithm was effective at capturing unfavorable genomic regions. The proportion of highly unfavorable ROH genotypes (i.e. < 0.05 percentile) that the algorithm captured under the high LD scenario varied across QTL scenarios. As the number of QTL increased the proportion of ROH genotypes captured decreased. The average (95% CI) proportion of highly unfavorable ROH genotypes the algorithm captured was 0.52 (0.42-0.62), 0.39 (0.32-0.46) and 0.33 (0.27-0.39) for the scenarios with 250, 500 and 1000 QTL, respectively. The prediction accuracy based on real data (average ± SD: 0.17±0.10) was roughly half of what was observed with the simulated data (high LD scenario mean (95% confidence interval (CI)): 0.49 (0.47-0.52)), although across the majority of traits, IIL had a prediction accuracy that was greater than zero. A prediction accuracy near zero was observed in LR for NBA and TNB, which may be due to multiple factors including purging of unfavorable ROH genotypes due to strong selection for initial litter size within the line as well as a smaller dataset than the one used in the LW population. In our study, whole genome regression based EGV resulted in a moderate predictive ability (average ± SD across trait and breed: 0.48±0.14). The use of a whole-genome regression method to benchmark the algorithm was used to illustrate the limitations of the algorithm. The algorithm only test for regions contained in longer ROH resulting in an unfavorable phenotype and should be used in conjunction with other methods to increase the overall genomic variability and limit the accumulation of inbreeding. Importantly, a moderate positive correlation between IIL and EGV was observed in the simulated (high LD scenario mean (95% confidence interval (CI)): 0.54 (0.53-0.56)) and swine (average ± SD: 0.31±0.10) datasets. Thus, the combination of the two metrics could allow for a breeder to more effectively manage the risks associated with sire or mate selection, allowing to better evaluate the trade-off between the genetic value of the progeny and undesirable side effects associated with inbreeding. Lastly, the use of the algorithm along with methods to identify lethal mutations/haplotypes (VanRaden et al., 2011) would allow breeders to comprehensively manage genomic diversity and recessive load in a population.

The two maternal lines utilized in this study have been under intense selection for multiple generations, which has potentially resulted in high and heterogeneous levels of homozygosity across the genome. This has been investigated recently by Howard et al. (2016), which estimated the proportion of the genome in a ROH of at least 5 Mb to be 0.17 and 0.19 for LR and LW, respectively. Furthermore, nearly all chromosomes across both breeds contained regions of the genome with high levels of ROH. Under this premise, it is likely that the impact of genome-level homozygosity would be regressed toward zero, since homozygosity in some regions of the genome would no longer be unfavorable. This result is partially verified by the ideogram outlined in **Figure S3** and **S4**, whereby some regions of the genome have unfavorable haplotypes spread across multiple traits and other do not have any unfavorable regions. The impact of haplotypes contained within an ROH for regions that were significant across multiple traits can be quite large. For example, an animal homozygous for a tag haplotype on SS9 (28.9-30.6) within the LR breed would be predicted to have 1.66 fewer pigs born alive, 1.32 fewer total pigs born, 4.0 % more pigs born dead and the litter would be on average 0.07 kg smaller than an animal not being homozygous for the tag haplotype.

In general, the genetic diversity of a population is managed through the relationship of the parents based on the expectation that the inbreeding in the progeny is equal to half of the coancestry between the parents (Falconer and Mackay, 1996). As previously discussed, since inbreeding depression is heterogeneous across the genome, a measure that has a higher relationship with the genetic load of an individual may serve as a better metric to manage the degree of inbreeding depression that exists within a population. Therefore, linear mixed models (i.e. Model 1 described in the section outlining the algorithm) that included either genome-wide inbreeding metrics or the IIL value in predicting a phenotype were evaluated and the corresponding *–log p-value* was estimated for each specific inbreeding regression coefficient. Across all simulated scenarios and on average across both swine breeds, the significance of the regression coefficient for the IIL value was higher compared to any other genome-wide metric. Yet for some traits genome-wide metrics were more significant. When both the most significant genome-wide inbreeding metric and ILL were included in the model, similar significance values remained. This highlight how genome-wide inbreeding and ILL metrics are capturing different signals. Based on our results, a combination of a genome-wide relationship matrix and **ILM** could be useful in effectively manage the risks associated with choosing an individual/mating combination. Future research should look at the long-term benefits of including the ILM in mating designs in terms of diversity and genetic load. Also, methods to incorporate multiple metrics including the genetic value, genetic diversity, lethal mutations and the unfavorable haplotypes from the algorithm into an index value should be developed.

Breeding objective are in the near totality of cases comprised of several economically important traits and thus the relationship between the **ILM** across traits is of importance. For the two breeds investigated the off-diagonal values across all traits resulted in a favorable or negligible change across all traits. Thus, the use of the **ILM** matrix for a given trait would result in a favorable increase or negligible change in the phenotype of the remaining traits. Furthermore, based on genome-wide relationship metrics (i.e. pedigree or genomic), the off-diagonals elements are favorably correlated with off-diagonal elements of the **ILM** matrix across traits and breeds. Therefore, as one changes the **ILM** values in the favorable direction, the relationship across mating pairs is reduced, which is desirable and expected. Future research should investigate methods to combine **ILM** across traits in the breeding objective. In general, the diagonal values had similar trends as the off-diagonals across traits and relationship matrices. One of the major differences between the two values related to an antagonistic relationship for LR between the SNPRM and the IIL values across all traits. The inbreeding correlations had a much lower correlation than off-diagonals elements and even more so within the LR breed.

In the present study variance components were not re-estimated for each window in Stage 2, which may have impacted the t-statistic. We utilized ASreml, which does re-estimate the variance components for each window, to determine the sensitivity of fixing variance components. The difference between the T-statistic from the algorithm and the one from ASReml is outlined in **Table S4**. Across all breeds and traits, the differences between the two were negligible. In addition, across all breeds and traits the T-statistic from the algorithm gave a conservative estimate compared to the ones derived from ASreml. The algorithm proposed may tag short haplotypes instead of long ones that aren’t most likely IBD segments as a result of recent inbreeding. Averaged across traits the proportion of ROH genotypes that were below 1, 2 and 3 Mb with LR (LW) was 2.0 (1.7), 8.3 (8.4) and 16.0 (16.2) percent, respectively. The algorithm trapped short ROH genotypes (albeit at a low frequency). Further improvements of the algorithm in the future should focus on reducing the frequency of trapping short ROH.

Previous studies have investigated ROH effects by accounting for the additive genotypic value of the region in the model by either including SNP contained within the region investigated (Pryce et al., 2014) or using phenotypes that have been corrected for the additive effect (Howard et al., 2015). When utilizing a separate model for each window, the simulation and swine data sets have illustrated that the ROH tagging haplotype can span many Mb and is variable across animals within and across windows. Therefore, the number of SNP to include before and after the haplotype in the model to account for the additive effect for a given region is difficult to determine. More importantly, the independence between additive and dominance effects in the classical treatment (Falconer and Mackay, 1996) is to an extent a convenient artifact that allows orthogonally of the additive and dominance estimates. In reality, and as outlined in Huang & Mackay (2016), depending on the parameterization of the model the variance explained by either additive, dominance or epistasis can be rearranged and placed more heavily into any of the three categories. This is chiefly due to the fact that three effects are in real situation non-orthogonal to each other, so that the variance from a particular effect can be “consumed” by another effect. This is an important point as additive and dominance are two intrinsically inseparable terms, since if an allele is dominant over another (a ≠ 0, d ± a), there must necessarily be additive homozygous effects (a ≠ 0; Huang and Mackay, 2016). The non-orthogonal relationship between additive and dominance effects has been confirmed with real data (Wellmann and Bennewitz, 2011; Wellmann and Bennewitz, 2012). The non-orthogonal relationship was also observed in the current simulation study. The correlation between ILL and true dominance deviation was essentially 0 across all scenarios, although a positive correlation (0.14) was observed between ILL and the estimated dominance deviation EDD was observed. Under this premise, the ability to efficiently estimate the additive and dominance effect and their potential interactions for QTL that are at a low frequency is severely reduced. Lastly, the application of the associated haplotypes identified in mating plans when correcting for the additive effect is even more complex due to a lack of clear interpretation between the combined additive and ROH effect for a window. Therefore, in our analysis priority was given to estimating the genotypic value of ROH segments that are susceptible to displaying reduced performance based on the combined genotypic value of the given segment. Based on this premise, we make no attempt at trying to understand the number of mutations present within the ROH, the degree of epistasis that occurs or the inheritance pattern of QTL within the segment.

### Conclusions

We have outlined an algorithm that identifies unfavorable haplotypes contained within an ROH that give rise to a reduced phenotype. Across simulated and real datasets the unfavorable haplotype tags a much larger ROH region that has a high probability of being IBD due to its length. Furthermore, the accuracy of prediction for the majority of the traits was greater than zero. On the real swine datasets, multiple haplotypes were identified that had a consistent unfavorable effect across multiple traits. The use of this algorithm and the associated haplotypes allow for breeding programs to more effectively identify unfavorable regions and mating programs can be used to minimize the frequency of ROH occurring in the next generation.

